# Identification of regional astrocyte heterogeneity associated with cuprizone-induced de- and remyelination using spatial transcriptomics

**DOI:** 10.1101/2024.03.04.583308

**Authors:** Anneke Miedema, Marion H.C. Wijering, Astrid M. Alsema, Emma Gerrits, Michel Meijer, Mirjam Koster, Evelyn M. Wesseling, Wia Baron, Bart J.L. Eggen, Susanne M. Kooistra

## Abstract

The cuprizone model is a well-characterized model to study processes of demyelination and remyelination, which are known features of multiple sclerosis. Cuprizone induces oligodendrocyte loss and severe demyelination in the brain, including the corpus callosum, hippocampus, and cortex. Loss of oligodendrocytes and myelin is accompanied by microgliosis and astrogliosis, wherein microglia and astrocytes partially lose their homeostatic functions and acquire a reactive/activated state. Cuprizone-induced demyelination peaks later in grey matter (GM) than in white matter (WM), and remyelination is more efficient in WM areas. Here, we aim to better understand regional diversity in microglia, astrocytes, and oligodendrocytes and their respective role in remyelination efficiency, by characterizing their response to cuprizone across brain regions. We applied spatial transcriptomics (ST) for unbiased gene activity profiling of multiple brain regions in a single tissue section, to identify region-associated changes in gene activity following cuprizone treatment. Gene activity changes were detected in highly abundant cell types, like neurons, oligodendrocytes, and astrocytes, but challenging to detect in low-abundant cell types such as microglia and oligodendrocyte precursor cells. ST revealed a significant increase in the expression of astrocyte markers *Clu*, *Slc1a3,* and *Gfap* during the demyelination phase in the WM fiber tract. In the cortex, the changes in GFAP expression were less prominent, both at the transcriptional and protein level. By mapping genes obtained from scRNAseq of FACS-sorted ACSA2-positive astrocytes onto the ST data, we observed astrocyte heterogeneity beyond the simple classification of WM– and GM-astrocytes in both control and cuprizone-treated mice. In the future, the characterization of these regional astrocyte populations could aid the development of novel strategies to halt the progression of demyelination and support remyelination.

**Highlights:**

⍰ Astrocyte markers *Clu*, *Slc1a3,* and *Gfap* are increased in WM fiber tracts during demyelination
⍰ Expression dynamics of astrogliosis markers *Gfap* and *Vim* during de-and remyelination depend on the brain region
⍰ Combining scRNAseq with ST data revealed astrocyte heterogeneity beyond WM– and GM-differences
⍰ scRNAseq-identified gene sets were differently affected by cuprizone treatment across brain regions

## Introduction

Multiple sclerosis (MS) is an immune-mediated neurodegenerative disease of the central nervous system (CNS). Hallmarks of MS include the presence of demyelinated lesions, inflammation, astrogliosis, and microgliosis (Lucchinetti et al., 2000; Schirmer et al., 2021). Microglia and astrocytes are important regulators of brain homeostasis and studies reported different roles of these cell types in MS progression (Guerrero & Sicotte, 2020; Ponath et al., 2018; Yong, 2022). Inflammatory processes induced by the activation of microglia and reactivity of astrocytes can be beneficial in MS, contributing to brain homeostasis via the control of iron metabolism and by providing neurotrophic support (Waller et al., 2016; Yong, 2022). However, during MS-related inflammation, microglia and astrocytes release substances that promote inflammation and neurotoxicity, leading to tissue damage and hindering the remyelination process by inhibiting the differentiation of oligodendrocyte-precursor cells (OPCs) (Traiffort et al., 2020).

Previous studies have identified regional heterogeneity of glial cell types in the adult mouse brain (Bayraktar et al., 2020; Marques et al., 2016; Schirmer et al., 2021; Tan et al., 2020). Using single-cell RNA sequencing, distinct astrocyte subpopulations were identified in the cortex compared to the hippocampus, cerebellum, thalamus, and hypothalamus (Lee et al., 2022; Tan et al., 2020). Microglia also differed in their number, morphology, and molecular signature across mouse brain regions (Lee et al., 2022; Tan et al., 2020). In addition, OPCs and mature oligodendrocytes (OLs) showed functional differences in distinct brain regions (Lentferink et al., 2018). Grey matter (GM) neonatal rat OPCs were less mature and possessed a higher proliferating capacity compared to white matter (WM) OPCs (Lentferink et al., 2018). Differences between regional glial subpopulations may lead to various responses of these subpopulations to pathological stimuli, highlighting the importance of understanding regional heterogeneity in the context of MS (Werkman et al., 2021).

Current studies on glial cell heterogeneity in MS or MS mouse models usually investigate one selected brain region, such as the spinal cord, cortex, or cerebellum (Bayraktar et al., 2020; Trobisch et al., 2022). Advancements in molecular technologies have recently enabled us to study gene expression profiles within a tissue section that contains multiple different brain regions (Ståhl et al., 2016). Ståhl et al. were the first to add this spatial component to RNA-sequencing technologies in 2016 at a spot resolution of 100 µm with spatial transcriptomics (ST). ST has contributed to a better understanding of the regional molecular changes in experimental autoimmune encephalitis (EAE), Alzheimer’s disease (AD), and amyotrophic lateral sclerosis (ALS) mouse models (Chen et al., 2020; Gadani et al., 2023; Maniatis et al., 2019), however, no demyelination mouse models have been studied so far using ST.

The cuprizone mouse model is a well-established model to study demyelination and successful remyelination processes (Leo & Kipp, 2022). Cuprizone is a copper-chelating reagent causing oligodendrocyte cell death in the mouse brain, resulting in demyelination. Demyelination is induced from the start of cuprizone treatment and is associated with astrogliosis and microgliosis (Gudi et al., 2014). Utilizing immunohistochemistry, demyelination and astrocyte reactivity are detectable from week 3 (Gudi et al., 2014). OPC proliferation and migration starts from week 3-4 after cuprizone induction. Subsequently, OPCs undergo differentiation into myelinating OLs at approximately week 5 (Gudi et al., 2014). When the cuprizone diet is discontinued, after week 5, this allows for the completion of the spontaneous remyelination process (Gudi et al., 2014; Leo & Kipp, 2022). Astrocyte responses to cuprizone-induced injury can hinder or promote remyelination by modifying the environment for oligodendrocytes and microglia-astrocyte crosstalk (Gorter & Baron, 2022), indicating that glial interactions play a crucial role in this process.

In this study, we used brain tissue sections derived from the cuprizone mouse model on the ST platform. Our goals were to 1) identify alterations in gene expression during de– and remyelination, 2) investigate the regional gene expression changes of glia cell type markers during de– and remyelination, and 3) examine if brain regions are differently affected by cuprizone treatment.

## Materials and methods

### Animals

C57BL/6J-Cx3cr1tm2.1(cre/ERT2)Litt Gt(ROSA)26Sortm14(CAG-tdTomato)Hze mice were used for all experiments. These mice were bred in-house on a C57BL/6J background. All mice in the study should express one copy of the tomato reporter since the breeding pair expressed the gene in a homozygous manner. Genotypes of the offspring were verified using PCR on a randomized basis. All animal experiments were approved by the national central authority for scientific procedures on animals (CCD) and performed following ethical regulations (Permit# AVD105002015360). To activate cre-recombinase and express the tomato reporter in Cx3cr1 expressing cells, 6-week-old animals were treated twice with 500 mg/kg body weight tamoxifen (Sigma-Aldrich, cat# T5648-5G) dissolved in corn oil (Sigma-Aldrich, cat# C8267-500ML), administered via oral gavage.

### Cuprizone mouse model

Cuprizone-induced oligodendrocyte intoxication was applied temporarily to the mice, which models the process of demyelination and successful remyelination. Demyelination was induced in 8-week-old male mice via a 0.2% w/w cuprizone diet (Sigma-Aldrich, cat# C9012-25G). The diet was freshly prepared by mixing cuprizone with standard powder food and water, and stored for up to one week at –20 °C. Animals were fed with this homemade chow three times a week, which they could eat ad libitum. Control animals received chow prepared similarly, lacking cuprizone. The experimental groups were early demyelination (3-week cuprizone diet), complete demyelination (5-week cuprizone diet), and remyelination (2-week withdrawal of the cuprizone diet). Brains were collected under deep anesthesia (4% isoflurane with 7.5% O2) after transcardial perfusion with 20 mL PBS. Experiments were performed in two batches. The mouse brains of the first batch were further processed for ST (n = 2) and astrocyte immunohistochemistry (n = 3). Subsequently, the brains of the second batch were used for acute astrocyte isolation and single-cell sequencing, where in one sample, the brains of 5 mice were pooled.

### Spatial transcriptomics

Directly after collection, brains were sectioned at bregma +1 and –3 and along the coronal plane, after which half of the brain was placed in a 10 by 10 mm plastic mold filled with OCT (Tissue-Tek, cat. no. 4583). Subsequently, the dissected mouse brain regions were fresh-frozen using cold isopentane (2-methylbutane; Fisher Scientific, cat. no. 10542331) and stored at –80C. This procedure was performed as quickly as possible to maintain tissue morphology and RNA integrity. To generate spatial gene expression profiles, the OCT-embedded mouse brains, ST glass slides, and a clean blade were pre-cooled in the cryostat chamber for 30 minutes. Subsequently, 16 µm thick cryosections were attached to the capture areas of an ST microarray slide. The size of a coronal section of one mouse brain hemisphere exactly fits the capture area of 6,2 by 6,6 mm. We cut the mouse brains in coronal sections between bregma –0.5 and –2 since most cuprizone-induced damage in the corpus callosum is expected in this region (Steelman et al., 2012). Each slide contains up to 6 sub-arrays, filled with 1007 spatial spots containing barcoded oligo dT probes. The spatial spots have a diameter of 100 µm and a spot-to-spot distance of 200 µm. ST was performed according to the protocol described previously (Salmén et al., 2018). Briefly, sections were post-fixed with 4% formaldehyde for 10 minutes and histologically stained with Dako Mayers hematoxylin (Agilent, cat. no. S3309) (4 minutes) and eosin Y (Sigma-Aldrich, cat. no. HT110216). The HE-stained tissues were imaged in black and white with a confocal microscope (Zeiss Cell Discoverer 7) and in color using a high-content fluorescence Widefield microscope (TissueFaxs). A scratch was made on the slide with a diamond pen to facilitate image alignment with spatial spots. Pre-permeabilization was performed with collagenase I (Gibco; Thermo Scientific, cat. no. 17018-029) for 20 minutes. Followed by permeabilization with pepsin (Sigma-Aldrich, cat. no. P7000-25G) for 10 minutes. Note; permeabilization enzyme and duration were optimized for brain tissue in advance using a tissue optimization slide. During permeabilization, the poly-A tail of the RNA molecules attaches to the oligo dT capture site. These probes also contain a unique spatial barcode, which is crucial for retaining spatial information. All pipetting steps in this protocol with RNA were performed in a flow cabinet that was cleaned with DNA and RNAse away, to protect the sample from contamination and keep the RNA intact. Subsequently, superscript III (Invitrogen; Thermo Fisher, cat. no. 18080085) was used for reverse transcription of the RNA overnight. The next day tissues were removed from the slide using proteinase K (Qiagen, cat. no. 19131) and cDNA was released from the glass slides and collected in low-binding tubes for cDNA purification and second-strand synthesis. The spatial spots on the slides were visualized with a Cy3 fluorophore. This fluorescent signal was imaged using the confocal microscope (Zeiss Cell Discoverer 7). Fluorescent and bright field images were aligned using Adobe Photoshop. Next, the cDNA was in vitro transcribed to RNA, amplified, and purified and RNA adapters were attached. Amplified RNA was transcribed to cDNA again for further construction of a sequencing library. Based on qPCR we proceeded with the indexing PCR for 10 cycles for slide 1 and 7 cycles for slide 2. After indexing PCR, the libraries were purified and concentrations were quantified using Qubit fluorometer (Invitrogen, cat. no. 33216) with a dsDNA HS Assay Kit (Invitrogen, cat. no. Q32854) and tapestation (Agilent 2200 TapeStation system) with high sensitivity kit (High Sensitivity D5000 ScreenTape, 5067-5592) and (High Sensitivity D5000 Reagents, 5067-5593). 1.1 pM of the library was loaded on an Illumina NextSeq 500 Sequencing System in the UMCG sequencing facility, and run using a NextSeq 500/550 High Output Kit v2.5 with 75 cycles (Illumina, 20024906).

### Immunohistochemistry

Mice from the same batch were used either for ST or for immunohistochemistry. 4% PFA-fixed frozen mouse brains were sectioned at a thickness of 16 µm between bregma –0.5 and 2. This region was selected since most cuprizone-induced demyelination is expected from bregma –0.5 (Steelman et al., 2012). Sections were stained using immunohistochemistry. For this method, heat-induced antigen retrieval with sodium citrate (pH = 6) was applied to unmask the epitopes. Next, sections were washed with PBS and incubated with 0.3% H2O2/PBS to block endogenous peroxidases for 30 minutes. Fc receptor blocking was performed for 1 hour using 5% normal goat serum (GFAP staining) or 5% horse serum (MOG staining) dissolved in PBS with 0.3% Triton X-100. After washing with PBS, sections were incubated overnight at 4 °C with the primary antibody: GFAP (Agilent/Dako, z033420-2, dilution 1:750) or MOG (Sigma-Aldrich, AMAB91067, dilution 1:1000) in PBS containing 1% normal goat serum (GFAP staining) or 1% horse serum (MOG staining) and 0.3% Triton X-100. The next day, sections were washed with PBS and incubated with a biotinylated secondary antibody: goat-anti-rabbit (Vector, BA1000, dilution 1:400) or horse-anti-mouse (Vector, BA2001, dilution 1:200), followed by a 30-minute incubation with the ABC solution (Vectastain elite kit, PK-6100). After rinsing with PBS, the sections were incubated for 10 minutes in DAB and 0.03% H_2_O_2_ (GFAP staining) or 5 minutes in DAB and 1.5% H_2_O_2_ (MOG staining). Finally, the sections were dehydrated and mounted with DepeX (Serva, 18243). Images were acquired using a NanoZoomer 2.0-HT Digital slide scanner C9600 (Hamamatsu Photonics). For quantification, per animal a 40x zoomed image of the cortex and corpus callosum (1 image per region) was converted to grayscale (8-bit images), and the percentage of GFAP positive pixels was measured by FIJI.

### Acute astrocyte isolation

Astrocytes were enzymatically isolated from fresh mouse brains (without cerebellum and olfactory bulb) (n = 5 mice per group). Brain tissues were mechanically dissociated on a glass slide on ice using a knife until a cell suspension was obtained. This was transferred to a tube containing enzyme solution with 2 mL PBS, 20 mg Protease from Bacillus licheniformis (Sigma Aldrich, cat# P5380-1G) and 20 µL L-cysteine, incubated on ice for 15 minutes while mixing every 5 minutes. After enzymatic dissociation, the single cell suspension was passed through a 100 µm cell strainer (Corning, cat# 21008-950), filled up with 15 mL enriched HBSS. Cells were pelleted by centrifugation for 10 min, 300 RCF at 4 C. Myelin was removed by 24% Percoll (Fisher Scientific, cat# 17-0891-01) and PBS density gradient centrifugation for 20 min, 950 RCF at 4 C, the pellet contains a mixture of glial cells. The cells were passed through a 35-μm nylon mesh, collected in round bottom tubes (Corning, cat# 352235), and sorted using a Beckman Coulter MoFloAstrios cell sorter. Fc receptors were blocked using cd16/cd32 monoclonal antibody (eBioscience, USA) for 10 minutes on ice. Subsequently, cells were incubated for 20 min on ice with anti-mouse ACSA2-APC (Miltenyi Biotec, USA). Astrocytes were sorted based on ACSA2 positive, DAPI negative, and tomato negative signals. The gating on DAPI was used to exclude damaged cells and on tomato to exclude Cx3cr1-expressing cells. 30,000 sorted cells derived from 5 mice were pooled in 5 µL HBSS without phenol red and directly loaded on a chip for single-cell RNA sequencing (scRNA-Seq).

### Single-cell RNA sequencing of astrocytes

The 10X Genomics platform was used for scRNAseq, performed according to the Chromium single cell 3’ reagent kit v2 user guide. Approximately 30,000 sorted cells were transferred to a microfluidic chip derived from the Chromium single cell A Chip Kit (10X Genomics, cat# 120236). Additionally, the chip was filled with a master mix for cDNA synthesis, partitioning oil, and gel beads. Cell lysis and cDNA synthesis occurred within the GEMs and subsequently, GEMs were lysed using a recovery agent followed by library preparation using a Chromium single cell 3’ Library & Gel Bead Kit v2 (10X Genomics, cat# 120237). cDNA amplification required 12 and indexing required 14 PCR cycles. Samples were sequenced on an Illumina NextSeq 500 Sequencing System with NextSeq 500/550 High Output Kit v2.5 (75 cycles) (Illumina, cat# 20024906). We loaded 1.8 pM of the library with 5% PhiX. 27 base pairs (bp) were used for read 1 and 56 bp for read 2 with indexes consisting of 9 bp.

### Data processing (spatial transcriptomics)

Using transparency channels, the bright field stained images (HE) and fluorescent images (Cy3) were aligned using Adobe Photoshop CS6. Using the open-source ST Pipeline (Navarro et al., 2017), the raw sequencing reads were processed with default settings. Reads were aligned to the genome reference Ensembl GRCm38 and reference Mouse GenCode release M31. Count matrices were filtered to contain only protein-coding, long-non-coding-intergenic, and antisense genes. Per the sample, the next filtering was to keep only those spatial spots located inside the tissue using the file generated by image alignment. The count data of all sections was merged into one Seurat object. In terms of quality criteria, spatial spots containing at least 100 features and less than 35% mitochondrial genes were included in downstream analysis.

### Dimensionality reduction and clustering (spatial transcriptomics)

The filtered expression matrix was processed using Seurat with default parameters (v.4.0.4)(Hao et al., 2021). Log-normalization was first performed and followed by FindVariableFeatures using selection.method=”vst” and default settings. Data was scaled and centered using the function ScaleData(). Principal component analysis was performed using RunPCA() calculating the top 50 dimensions. Next, clustering was performed using functions FindNeighbors(), FindClusters(), and RunUMAP(), using dimensions 1:30 and a resolution of 0.4 in addition to default parameters.

### Manual region annotation (spatial transcriptomics)

A custom R shiny app was developed to annotate each spot manually with the labels “cortex”, “WM fiber tract” and “hippocampus” based on the HE images. The WM fiber tract was defined as corpus callosum plus the myelin fiber tracts in other regions. The remaining spots were annotated as “not specified”.

### Region gene set annotation (spatial transcriptomics)

Raw counts were log-normalized using Seurat. A list of region-specific gene sets for 9 different brain regions was retrieved from Lein et al., 2007 (Table S2). We manually selected genes that are known to be enriched in WM fiber tracts (provided in Table S2). Next, we applied the AddModuleScore function from Seurat per region gene set and plotted the module score. Observations with a module score > 0.1 were considered as having a positive module score. Spots without a positive module score were annotated as “unidentified”. Some ambiguous spots had positive scores for multiple gene sets, likely reflecting spots on the border of anatomical brain regions. In this scenario, we annotated the spot to the region with the highest module score.

### Combinatorial approach for region annotation (spatial transcriptomics)

We initiated the combinatorial approach by comparing three methods for region annotation, namely manual annotation, clustering, and region genesets. This comparison was specifically conducted for the WM fiber tract, as this region exhibits the most easily identifiable cuprizone-induced injury. The comparison indicated that the manual annotation and clustering methods exhibited the highest level of agreement in annotating spots to the WM fiber tract. Spots were assigned to the WM fiber tract if they were annotated as such by either the manual or the clustering method (utilizing a full outer join of the Venn diagram). This means that the combinatorial approach accounted for both methods and encompassed the union of their respective results. The same combinatorial approach was applied for the hippocampus and cortex category. Ambiguous spots at the border of brain regions, where manual and clustering methods differ, were analyzed twice for region averages: once as belonging to the manually annotated region and once according to the clustering region. The combinatorial approach was used to calculate the mean expression of a gene/gene set per region or for pseudobulk analysis.

### Pseudobulking (spatial transcriptomics)

Per sample, we extracted the spatial spots that were likely to reflect a gene expression profile from WM fibers using the combinatorial approach for region annotation. The extracted spatial spots were either annotated by eye to be located within WM fibers or by clustering defined to be part of the WM fiber cluster. Next, we summed the UMI counts of the spots belonging to a given sample creating a pseudobulk. Principal component analyses were computed on variance-stabilizing-transformed pseudobulked count data using the ‘prcomp’ function. The input for PCA consisted of the top 2000 most variable genes.

### Differential expression analysis (spatial transcriptomics)

Genes were removed if they were not expressed in at least 5% of the spots. To compute differentially expressed genes in WM fibers between cuprizone and control mice, while accounting for sample– and slide-related variability, we applied linear mixed models with the formula: ∼ group + (1|sample_id) + (1|slide_number) using the R package variancePartition (v1.20.0) (Hoffman & Roussos, 2021). The count and metadata information for the model were provided on the spot level. “Group” represented the cuprizone treatment condition. Control mice (n = 2) were set as the reference group. Hypothesis tests were performed with t-tests based on Satterthwaite’s method. P-values were adjusted for multiple comparisons using the Benjamini & Hochberg method. Genes were defined as differentially expressed when the log2 fold change was > 0.5 or < –0.5 and the adjusted p-value was < 0.05. This log2 fold change threshold was chosen as an intermediate value between the conventional threshold used in bulk RNA sequencing (>1) and scRNAseq (>0.25).

### Gene set enrichment (spatial transcriptomics)

The differentially enriched genes, defined as changes with a log2 fold change > 0.5 and adjusted p-value < 0.05 in the pairwise comparisons D3-C, D5-C, R-C, D5-D3, R-D5, were given as a list for gene set enrichment using the web tool: https://www.flyrnai.org/tools/single_cell/web/enrichment. “Mouse” was used as the selected input gene species and “Gene Symbol” as the gene identifier. The “Brain_mm_Rosenberg” was used as a reference data set (Rosenberg et al., 2018), and the top marker genes per cell type were set to “Top 100”. The enrichment p-value was based on a hypergeometric distribution (Hu et al., 2021) and illustrated using the negative log10 of the p-value.

### Module scores (spatial transcriptomics)

Raw counts were log-normalized using Seurat. We applied the AddModuleScore function per gene set (e.g. PANreactive astrocyte gene set, scRNAseq-derived astrocyte cluster marker sets, others). To construct a spatial plot per gene set module score ggplot2 was used. Identical procedures were applied for spatial transcriptomics data from ST and Visium platforms. PANreactive, neuroinflammatory, and neuroprotective astrocyte gene sets were derived from Liddelow et al., 2017 and are provided in Table S2.

### Data processing (single-cell RNA sequencing)

Raw reads were aligned to the GRCm38 genome for mouse samples, using Cell Ranger (v3.0.0) with default settings. Raw count files were loaded into R (v3.6) and barcode filtering was performed with thresholds at >600 unique molecular identifiers (UMIs) for mouse cells (Lun et al., 2019). The multiplet rate mentioned in the 10x Genomics User Guide was used to set an upper threshold per sample for the number of UMIs per cell and remove doublets. Additionally, 7 small clusters of contaminating, non-astrocyte cell types were identified and removed from the data. Cells with a mitochondrial content >5% were removed from the dataset. There was only one count file per sample (n = 1), but a sample represented pooled astrocytes from the brains of 5 mice. For the mouse dataset, count files from different conditions were merged into one and further analyzed with Seurat.

### Dimensionality reduction and clustering (single-cell RNA sequencing)

The data were normalized by dividing the counts of each gene by the total sum of counts per cell and multiplied by a scale factor of 10,000 and log-transformed. Highly variable genes (HVGs) were calculated using the FindVariableFeatures function with selection.method = vst. The data was scaled and confounding variation associated with the number of UMIs and mitochondrial and ribosomal content was regressed out. Principal component analysis was performed using Seurat RunPCA() calculating the top 20 dimensions. Single astrocytes were clustered using functions FindNeighbors(), FindClusters(), and RunUMAP(), using dimensions 1:18 and a resolution of 0.7 in addition to default parameters. Cluster markers were computed using FindAllMarkers with only.pos = TRUE, test.use = “MAST” in addition to default parameters. These cluster markers were used to plot cluster module scores in the ST data. Gene ontology (GO) analysis was performed with Metascape, v3.5.20230501, at the interactive web tool metascape.org using default settings.

### Differential abundance testing of astrocyte subpopulations (single-cell RNA sequencing)

To assess the differential abundance of astrocyte subpopulations in the single-cell data, we employed scCODA (v0.1.9), which utilizes a Bayesian framework for joint modeling of all measured cell-type proportions, as opposed to treating them individually (Büttner et al., 2021). The nature of our one-sample dataset, consisting of pooled astrocytes from the brains of 5 mice, rendered traditional frequentist tests infeasible. However, with the Bayesian approach offered by scCODA, it is possible that one-sample datasets can attain significance, in specific scenarios where very large increases (e.g. absolute increases of 2000 cells in case versus control) are measured in abundant cell types (Büttner et al., 2021). In our analysis, we designated astrocyte cluster 10 as the reference cell type. scCODA automatically selected this cluster due to its minimal total dispersion, indicating its suitability as a reference. The false-discovery rate (FDR) level was adjusted to 0.2 instead of the nominal FDR 0.05 to increase the sensitivity for the low-replicate scenario, however, at the cost of a higher false discovery rate. For this reason, at FDR 0.2 we did not interpret the scCODA-inferred ‘credible effects’ or conclude any significance, but we used the scCODA-inferred ‘Final parameter’ and ‘log2 fold changes’ to rank the largest differences in cluster abundance.

### Average module scores per brain region (spatial transcriptomics)

To generate a line graph representing the average expression of a gene or gene set per region, we used the region annotation obtained through the combinatorial approach (IV), as explained earlier. The gene expression for spatial spots in a given region was averaged per sample and depicted as individual data points. In addition, gene expression or module scores in individual spots for a given region (e.g. hippocampus) were averaged per condition (control, D3, D5, and R) and depicted as a horizontal line.

### Visium spatial transcriptomics

High-resolution spatial gene expression data and corresponding fluorescent images were obtained from https://www.10xgenomics.com/resources/datasets/adult-mouse-brain-section-1-coronal-stains-dapi-anti-neu-n-1-standard-1-1-0. This data derived from publicly available datasets provided by 10x Genomics. The fresh-frozen coronal adult mouse brain section was processed using the Spatial 3’ v1 chemistry and stained with anti-NeuN (indicated in red) and DAPI (indicated in blue). Module scores for astrocyte cluster gene sets were computed and visualized as described above for the other ST data.

## Results

### Characterization of demyelinated regions in cuprizone-treated mouse brains using spatial transcriptomics

To investigate gene expression changes related to successful de– and remyelination, we applied ST to an acute cuprizone mouse model. Mice were fed with a normal diet (C) or cuprizone for 3– (D3) or 5 weeks (D5) to induce demyelination, then followed by 2 weeks on a normal diet to allow for remyelination (R). Additional sample information is provided in Table S1. Coronal mouse brain sections of one hemisphere between bregma –0.5 and –2 were placed on the capture area of barcoded slides for spatial gene expression analysis (Fig. 1A). The spatial spots have a diameter of 100 µm and spot-to-spot distance of 200 µm, resulting in averaged gene expression profiles of approximately 10-30 cells per spot (Fig. 1A). The brain sections were stained with hematoxylin-eosin (HE) to provide spatial orientation in the tissue (Fig. 1B). In our spatial spot gene expression profiles, approximately 2000 genes and 4000 unique transcripts were detected per spot (Fig. S1A-B). These numbers of genes and transcripts were comparable to the original protocol (Salmén et al., 2018; Ståhl et al., 2016) that identified 1500 genes and 3000 unique transcripts per spot in the mouse olfactory bulb. In addition, we detected on average 15% of reads derived from mitochondrial genes and 4% of ribosomal genes, equally distributed across the experimental groups (Fig. S1C-D). Using ST, the detected expression levels of genes representative for OPCs, homeostatic microglia, and reactive microglia were low in our ST data (Fig. 1C), impeding the detection of processes related to these cell types such as remyelination. In contrast, we were able to successfully identify transcriptional markers for neurons, oligodendrocytes, and astrocytes (Fig. 1C).

**Figure 1.**
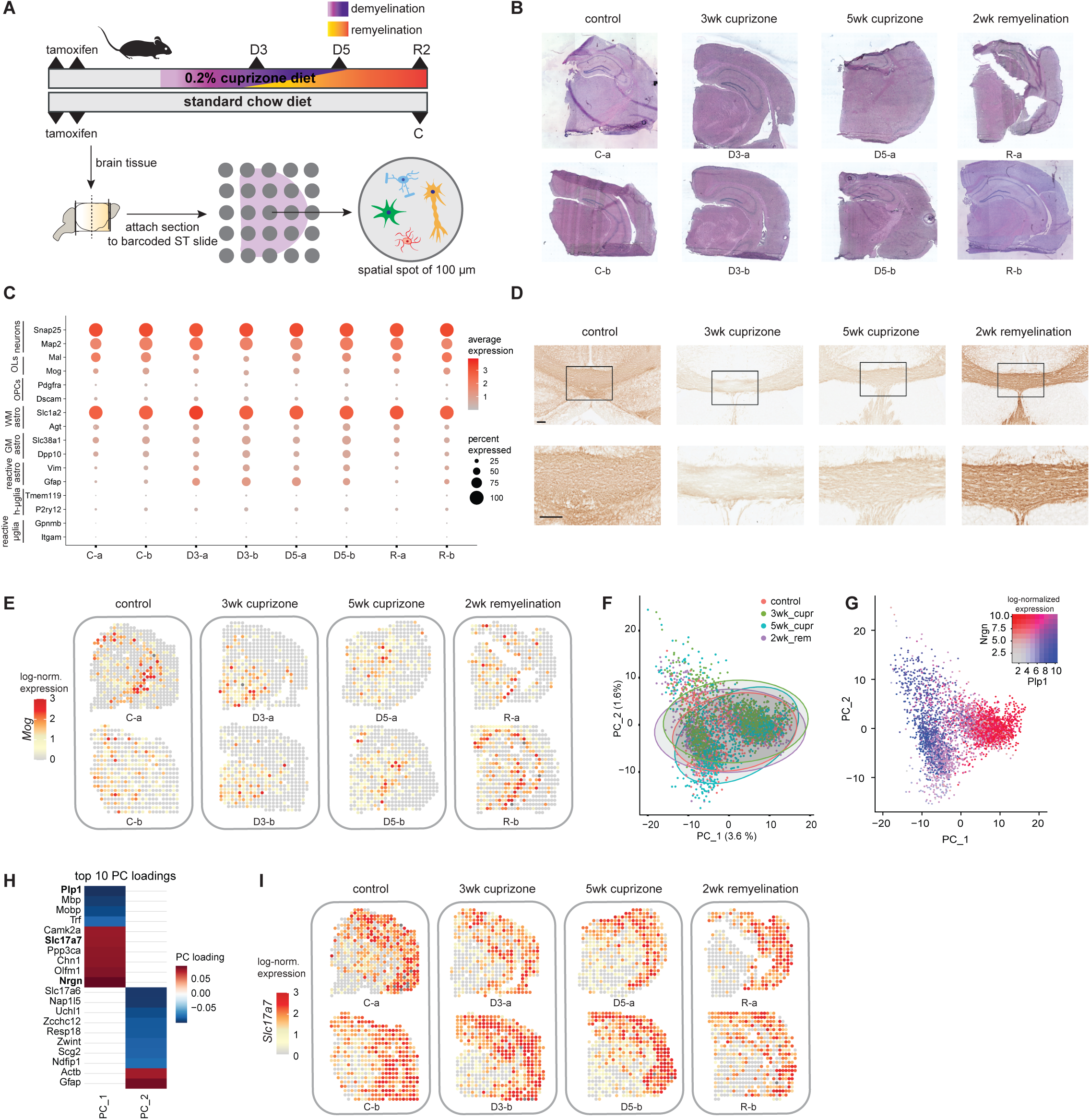
Detection of demyelinated areas in the brain of cuprizone-treated mice. **A**). Schematic overview of the cuprizone experiment. Mice received either a cuprizone diet to induce demyelination or a chow diet (control; C). At 3 (D3) or 5 weeks (D5) demyelination, or after a 2-week recovery period (R) following demyelination, mice were sacrificed and brains were collected. Brain samples were further processed for spatial transcriptomics (n = 2 per group). **B).** Hematoxylin and eosin (HE) immunohistochemistry of the ST brain hemispheres. **C).** Dot plot depicting the average expression levels of representative marker genes for neurons, oligodendrocytes (OLs), oligodendrocyte-precursors cells (OPCs), WM-, GM– and reactive astrocytes, and homeostatic (h-µglia) and reactive microglia per sample. The color scale depicts the average expression level. The size of the dots indicates the percentage of spatial spots expressing the gene. **D).** Immunohistochemical images of Mog expression in the corpus callosum (CC) per experimental group. The scale bar indicates 100 µm. **E).** Representative spatial gene expression plots of *Mog* per experimental group. The color scale indicates log-normalized gene expression levels. **F).** PCA plot of the spatial spots from all samples. Each dot represents a spatial spot, colors indicate experimental groups; control n = 1049 spatial spots; 3wk demyelination n = 990 spatial spots; 5wk demyelination n = 1004 spatial spots; 2wk remyelination n = 913 spatial spots. **G).** *Plp1* and *Nrgn* gene expression plotted on the PCA plot of the spatial spots. Each dot represents a spatial spot. The red color indicates the log-normalized expression of *Nrgn*, blue color indicates the log-normalized expression of *Plp1* in a spatial spot. **H).** Heatmap of the top 10 genes with the highest absolute PC loadings on PC1 and PC2. The color scale indicates positive versus negative PC loading. **I).** Representative spatial gene expression plots of *Slc17a7*. The color scale indicates log-normalized gene expression level. C = control, D3 = 3 weeks of cuprizone treatment, D5 = 5 weeks of cuprizone treatment, R = 2 weeks remyelination

First, we validated if the ST technique detected the decreased levels of myelin in the brains of cuprizone-fed mice. Demyelination in cuprizone-treated mice occurs throughout the whole brain, however, it is most readily detectable in the WM fiber tract at the protein level (Gudi et al., 2014). We detected a decreased level of the myelin oligodendrocyte glycoprotein (MOG) at 3 and 5 weeks of cuprizone diet, and MOG protein levels were increased after 2 weeks of remyelination (Fig. 1D). At the RNA level, a similar MOG expression pattern was observed (Fig. 1E).

Next, we performed principal component analysis (PCA) on all spatial spots to assess (dis)similarities in gene activity detected in ST spots. In the first principal components, spatial spots did not segregate based on the experimental group, but rather by white versus grey matter differences as indicated by *Plp1* and *Nrgn* expression levels in the PCA plot (Fig. 1F-G). The 10 genes with the highest absolute loadings for PC1 and PC2 were plotted to investigate the biological properties associated with these PCs (Fig. 1H). Expression of myelin genes, such as *Plp1* and *Mbp,* and in reverse direction neuronal-related genes, such as *Nrgn* and *Slc17a7*, had the highest loading on PC1 (Fig. 1H). Neuronal genes were expected to be most abundant in GM areas, such as the cortex. By mapping genes to spatial locations, we detected that the expression of neuronal gene *Slc17a7* was indeed most abundant in the cortex and less abundant in WM regions (Fig. 1I). Genes with the highest loading on PC2 included *Gfap*, suggesting the presence of reactive astrocytes in a subset of spatial spots (Fig. 1H).

Taken together, using ST with 100 µm resolution we observed cuprizone-induced alterations in expression levels of the myelin gene *Mog,* which was validated on protein level. However, based on PCA, the strongest gene expression variation in the data was related to gene activity associated with WM and GM areas rather than the experimental group. This indicates that further evaluation of the cuprizone treatment requires a region-specific analysis of gene activity using an accurate annotation of anatomical regions.

### Methods for annotation of anatomical regions in the mouse brain

The mouse brain contains different anatomical regions with diversity in cell type composition. In addition to anatomical heterogeneity, extensive cellular heterogeneity exists within a brain region, for example, neurons in the multiple cortical layers or the dentate gyrus within the hippocampus (Batiuk et al., 2020; Bayraktar et al., 2020). We assessed four strategies for annotating anatomical regions, each method presenting different advantages and limitations. Accurate annotation of brain regions for spatial spots is crucial to compare the experimental groups and detect cuprizone-induced changes. The methods evaluated were manual annotation (I), unbiased clustering (II), region geneset set enrichment (III), and a combinatorial approach (IV) (Fig. 2A). For method I-III the spatial plots were depicted per region (Fig. 2B). In method I (manual), we determined the exact bregma point of our samples and compared the histological HE stainings (Fig. 1B) with the Allen Brain Reference Atlas. We manually annotated the spots corresponding to the hippocampus, cortex, and WM fiber tract (Fig. S2A). WM fiber tract was defined as the corpus callosum and myelin fiber tracts in other brain areas. These three regions were easily recognizable areas by eye based on HE-histology. In this method the anatomical annotations were not affected by gene expression changes in response to the cuprizone treatment. However, this annotation method was restricted to identifiable regions, and potential subjectivity introduced by the observer cannot be excluded. Method II was based on unbiased clustering of gene activity in spatial spots identifying 13 spatial clusters (Fig. 2B). However, some clusters could not be clearly identified in all samples, such as the WM fiber tract-associated cluster 6. These findings suggest that clustering primarily results from regional variation but is also influenced by treatment-induced alterations in gene activity, observed in the WM fiber tract (e.g., cluster 6 affected by the cuprizone diet) and by the bregma point (e.g., cluster 5, absent in some samples) (Fig. 2B). In method III, we spatially plotted region-specific gene sets retrieved from literature (Lein et al., 2007) for annotation (Fig. 2B). Method III was effective in the annotation of various distinct anatomical regions and assigned a substantial number of spatial spots to the WM fiber tract (Fig. 2B, S2B). However, by visual inspection the region geneset enrichment method tended to assign spatial spots within the cortex to the WM fiber tract, suggesting this method may be inaccurate for annotating the WM fiber tract (Fig. 2B). Indeed, in the context of annotating WM fiber tract, the unbiased clustering method had greater overlap with manual annotation method compared to the geneset enrichment method. This observation suggests that the clustering method is more accurate than the region gene set enrichment method to computationally annotate WM fiber tracts (Fig S2C). Summarizing, each method had inherent advantages and limitations, but all methods added information on the brain region that corresponded to the separation of spatial spots in principal components 1 and 2 (Fig. S2B).

**Figure 2.**
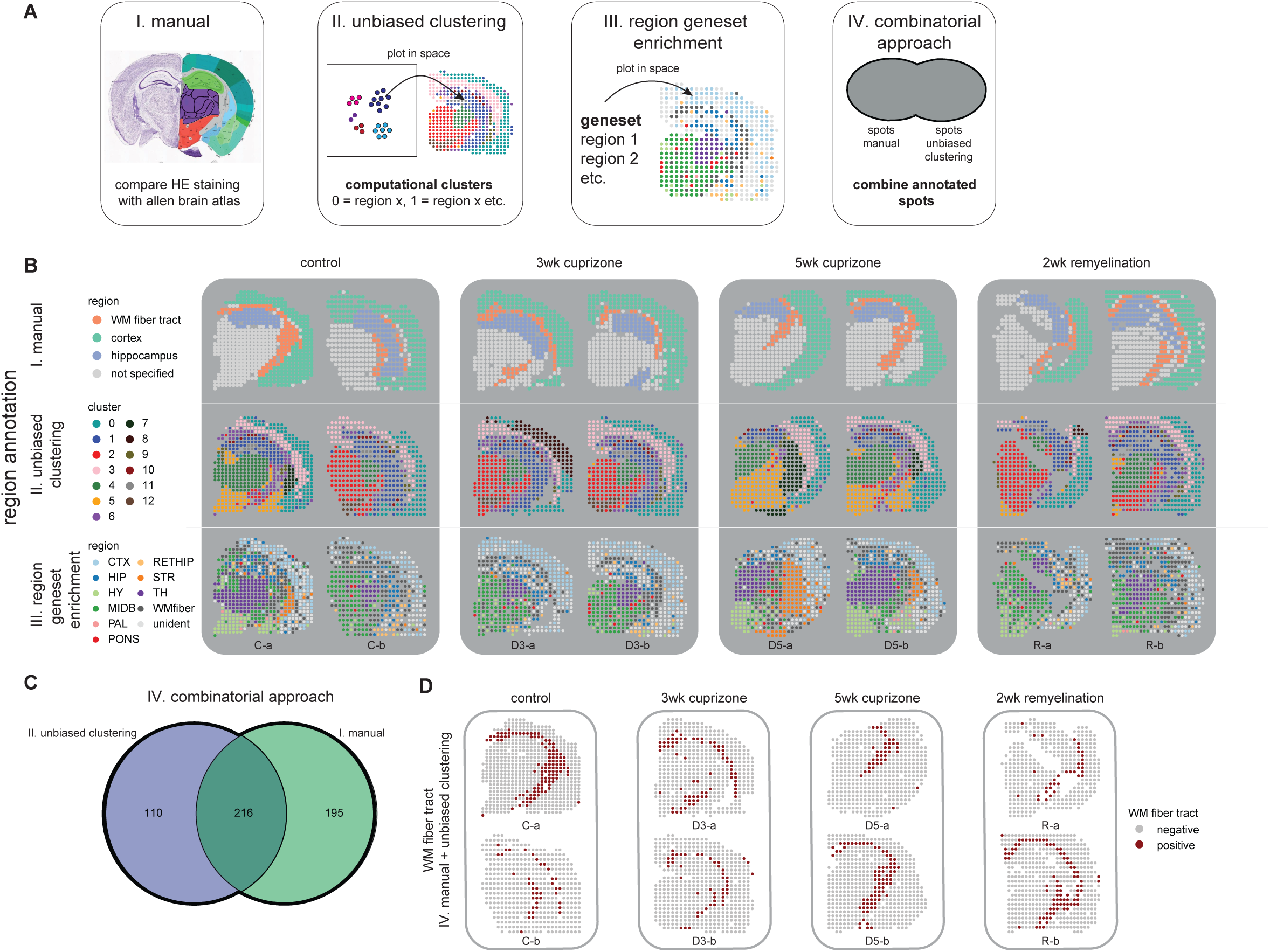
Comparison of different methods to annotate mouse brain regions. **A**). Schematic overview of four different methods to annotate brain regions. I: manual method; the Allen Brain Atlas was used to make an overlay between the HE-stained ST section and the atlas image. Spatial spots were manually annotated by comparing the section to the atlas. II: unbiased clustering method; unbiased clustering was performed on all spatial spots. By comparing each cluster’s location to the Allen Brain Atlas, clusters were annotated with a brain region. III: region geneset method; based on Lein et al., 2007, 10 region-specific gene sets were used to compute a region module score per spatial spot. Spots that expressed a gene module above a certain threshold were annotated to the corresponding region. IV: combinatorial approach of method I and II by using the full outer join of the spots positive for the manual and unbiased clustering method. **B).** Representative spatial plots of the different region annotation methods. Colors indicate different clusters or brain regions. **C).** Schematic overview of the combinatorial approach for WM fiber tract spot annotation. The manual method (I) and unbiased clustering method (II) are depicted as a Venn diagram. Spots annotated by either or both methods, shown as a bold line in the diagram, were annotated as WM fiber tract (n = 521 spots). **D).** Representative spatial spots for the WM fiber tract as a result of the combinatorial approach (IV). Dark red spots indicate the spots annotated to the WM fiber tract region. C = control, D3 = 3 weeks of cuprizone treatment, D5 = 5 weeks of cuprizone treatment, R = 2 weeks remyelination, CTX = cortex, HIP = hippocampus, HY = hypothalamus, MDB = midbrain, PAL = pallidum, RETHIP = retrohippocampal formation, STR = striatum, TH = thalamus, WM fiber = WM fiber tract, unident = unidentified.

To combine the most comprehensive manual annotation of the WM fiber tract with the most accurate computation method, we applied a combinatorial approach (method IV) wherein positive spots of either method I (195 positive spots) and method II (110 positive spots) or double positive spots (216 spots) were annotated to the specific brain region (Fig. 2D). This method gave a reliable annotation of the WM fiber tract, hippocampus, and cortex in cuprizone and control mice and allowed for a region-specific analysis of gene activity changes in this study (Fig. 2D, Fig. S2D-E).

### Gene expression dynamics of glia markers in the cuprizone mouse model

Using a combinatorial approach (method IV), we could distinguish gene activity in the WM fiber tract, hippocampus, and cortex, which was no longer masked by regional heterogeneity (Fig. S3A). Per region, we investigated the average expression levels of myelin-related, microglial, and reactive astrocyte genes at different time points of cuprizone treatment.

Based on previous literature, *Pdgfra* and *Dscam* were considered OPC markers (Huang et al., 2020; Marques et al., 2016). Detection of OPC markers was limited in ST data, making it challenging to investigate the dynamics of OPC-related gene expression per region using ST (Fig. S3A). *Opalin,* also known as *Tmem10,* is a marker of myelinating OLs and is highly upregulated during the early stages of OPC differentiation (de Faria et al., 2019; Marques et al., 2016). In contrast to other OPC markers, for this gene, some expression signal was detected, with a slightly higher expression observed in the R group than in the D3 group within the WM fiber tract (Fig. S3A). This could indicate two scenarios, either the relative abundance of other cell types is decreased, leading to an increase in detected *Opalin* transcripts. Alternatively, there might be a higher proportion of OPCs in the WM fiber tract at the remyelination phase, contributing to the observed increase in *Opalin* detection.

Expression of *Mal*, a marker specific to mature myelinating oligodendrocytes (Kuhn et al., 2019; Marques et al., 2016), was decreased in D3 compared to C mice and partially returned to control expression levels at D5 in the WM fiber tract and returned to control expression levels at R within the cortex and hippocampus (Fig. S3B). *Plp1* and *Mbp* are expressed at multiple stages during OL differentiation (Kuhn et al., 2019). *Mbp* is expressed in immature and myelinating OLs, while *Plp1* is expressed in OPCs, immature OLs, and also in myelinating OLs (Kuhn et al., 2019). *Plp1* and *Mbp* had similar gene expression patterns as *Mal* during cuprizone treatment (Fig. S3B). The incomplete reduction of these myelin markers at D5 can be explained by the concurrent processes of de– and remyelination that occur between D3 and D5 (Gudi et al., 2014). Expression of microglial genes such as *Tmem119* and *Itgam* was detected at very low levels in the ST data, hindering the examination of differences in microglia gene expression at different time points (Fig. 1C, Fig. S3C).

In contrast, a clear increase in the expression of reactive astrocyte genes *Gfap* and *Vim* was observed at D3 in the WM fiber tract and hippocampus compared to controls (Fig. S3D). The peak in gene expression for *Gfap* and *Vim* was earlier in the WM fiber tract (D3) than in the cortex (D5) (Fig. S3D). Following this peak in expression at D3, expression levels of *Gfap* decreased again from D5 in the WM fiber tract and hippocampus but did not return to control levels. For *Vim,* the decrease in gene expression after D3 varied per brain region (Fig. S3D). These results suggest that the expression of genes related to astrogliosis peaks at distinct time points for different brain regions.

### Cuprizone treatment increases the expression of reactive astrocyte-related genes in the WM fiber tract

Given the observation that alterations in selected glia markers were most noticeable in the WM fiber tract, we aimed to perform an unbiased differential expression analysis (DEA) within the WM fiber tract following cuprizone treatment. PCA indicated that the variation in spatial gene expression profiles within the WM fiber tract was associated with the different experimental conditions. For example, control mice separated from mice treated with cuprizone for 3 weeks (D3) (Fig. 3A). Within the WM fiber tract, we detected in total 70 differentially expressed genes (DEGs) in 5 comparisons (D3-C, D5-C, R-C, D5-D3, R-D5) (Table S3) and two comparisons (D3-C, R-D5) were summarised in the four-way plot (Fig. 3B). Gene set enrichment of the DEGs within the WM fiber tract indicated that mostly astrocyte-related DEGs, and few neuronal and myeloid cell-related DEGs were detected (Fig 3C, Table S4). DEGs identified between control and cuprizone-fed animals included astrocyte genes such as *Clu* and *Slc1a3* (Fig. 3D, Table S3). *Clu* expression was significantly enriched in both D3 and R versus control WM fiber tracts (Table S3). *Slc1a3* was enriched in the D3 versus control WM fiber tracts (Table S3). To get insight into the role of these astrocytes in de– and remyelination, gene sets of known pathological astrocyte subpopulations were plotted onto our spatial data. No differences in the gene expression of markers associated with inflammatory (e.g. *Ggta1, Fbln5*, etc.) (Liddelow et al., 2017) or neuroprotective astrocytes (e.g. *Clcf1, S100A10,* etc.) (Liddelow et al., 2017) were observed in the WM fiber tract of cuprizone treated mice (Fig. S4A-B). A PANreactive astrocyte gene set (*Lcn2, Steap4, S1pr3, Timp1, Hspb1, Cxcl10, Cd44, Osmr, Cp, Serpina3n, Aspg, Vim,* and *Gfap*) (Liddelow et al., 2017) was abundantly expressed within the WM fiber tract of cuprizone treated mice and seemed enriched upon cuprizone treatment (Fig. 3E). Increased *Gfap* and *Vim* expression was observed upon cuprizone treatment in the WM fiber tract and hippocampus (Fig. 3F, Fig. S3D and S4C). In comparison, the astrocyte marker gene *Aldh1l1* (Cahoy et al., 2008) was expressed throughout the whole mouse brain (Fig. S4D). This indicates that cuprizone induces a reactive gene set in WM fiber tract astrocytes during demyelination.

**Figure 3.**
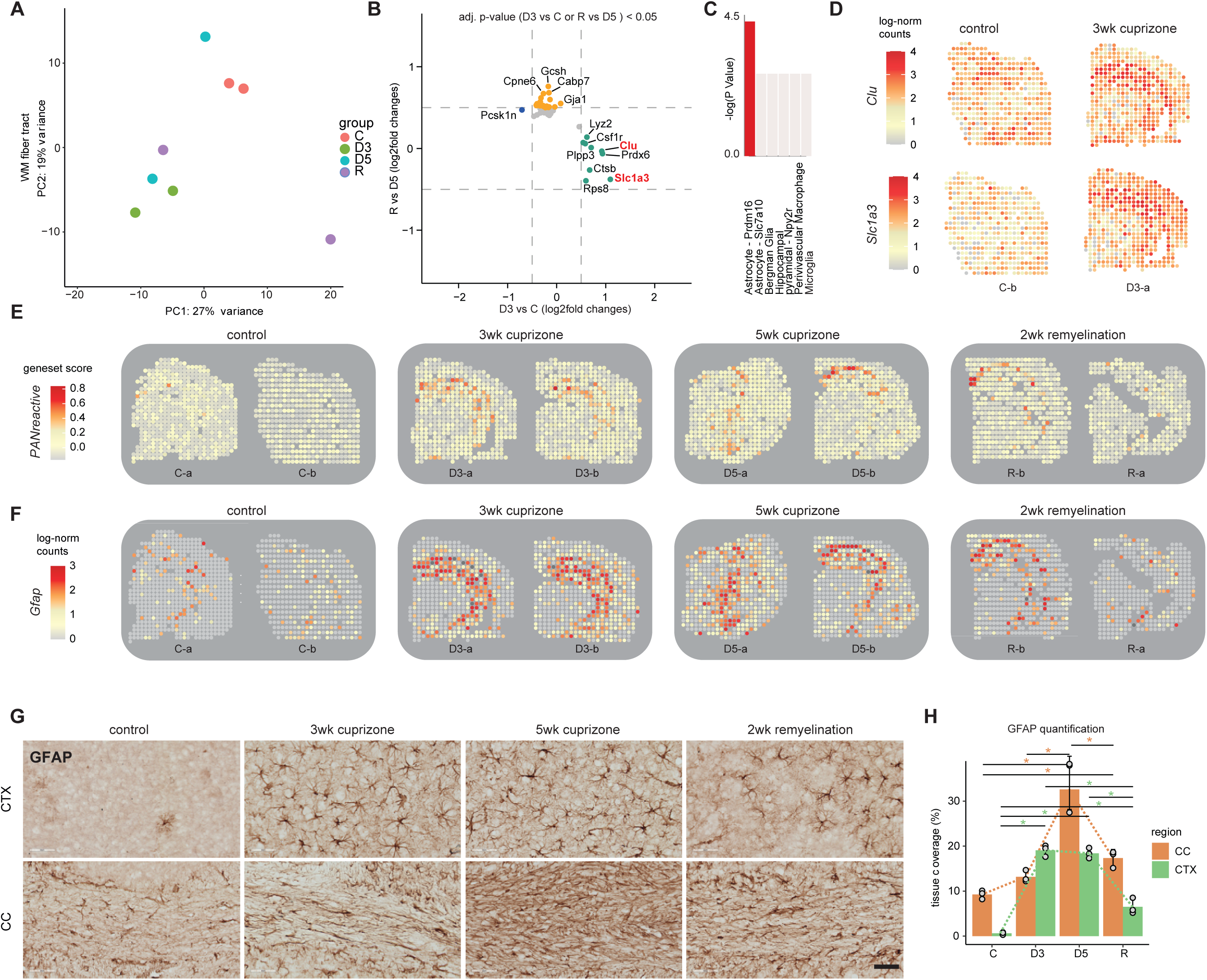
Differences in gene expression between cuprizone and control WM fiber tracts. **A**). PCA plot of pseudobulked WM fiber tract spots per experimental group. Each dot represents a sample of the WM fiber tract, colors indicate the experimental groups. **B).** Four-way plot comparing differential gene expression observed in WM fiber tracts of control and cuprizone-treated mice. Each dot represents a gene that is significantly changed in its expression. Green and blue dots are genes significantly increased or decreased in expression in D3 compared to C mice. Yellow dots represent genes significantly increased in expression in R compared to D5. We did not detect genes significantly decreased in expression in R compared to D5. Genes with a red label are visualized in spatial plots in d **C**). Bar plot depicting gene set enrichment of differentially enriched genes compared to a reference data set of mouse brain cell types (Rosenberg et al., 2018), computed using DRscDB (Hu et al., 2021). The height and color of the bar indicate the negative log10 of the enrichment p-value. **D**). Representative spatial gene expression plots of astrocyte genes *Clu* and *Slc1a2*. The color scale indicates log-normalized gene expression level. **E).** Representative spatial gene expression plots of PANreactive gene set scores. The color scale indicates the gene set module score. **F).** Representative spatial gene expression plots of *Gfap*. The color scale indicates log-normalized gene expression level. **G).** Immunohistochemical images of GFAP expression in cortex (CTX) and corpus callosum (CC). The scale bar indicates 50 µm. **H).** Quantification of immunohistochemistry of GFAP in cortex and corpus callosum shown in **e**. Per brain region, one-way ANOVA with Tukey HSD post-hoc tests were performed to assess the significance of GFAP levels among groups, n = 3 per group. Asterisks indicate p-values < 0.01. C = control, D3 = 3 weeks of cuprizone treatment, D5 = 5 weeks of cuprizone treatment, R = 2 weeks remyelination

Astrocyte reactivity in cuprizone mice was corroborated at the protein level. During cuprizone treatment, there were significant changes in the level of GFAP-positive astrocytes in both the cortex (CTX) and the WM fiber tract – corpus callosum (CC) (one-way ANOVA per region, n = 3 per group; CTX p<0.001, CC p<0.001), indicating local astrogliosis in these regions (Table S5, Fig. 3G-H). Astrogliosis persisted at the remyelination phase in the CTX and CC (Tukey HSD, n = 3 per group; CTX p<0.001, CC p = 0.005), suggesting a role for astrocytes in remyelination (Table S5, Fig. 3G-H). Generalized linear mixed models demonstrated a significant interaction between region and cuprizone treatment (p < 0.001, n = 3 per group) (Table S5, Fig. 3H), indicating that cuprizone treatment differentially affects astrogliosis depending on the brain region.

### Regional heterogeneity of astrocyte subpopulations in the cuprizone mouse model

To investigate if regional astrocyte diversity is associated with de– and remyelination, we combined scRNAseq of astrocytes with the spatial information on mouse brain regions. ACSA2, a known astrocyte marker (Batiuk et al., 2017; Borggrewe et al., 2021), was used to enrich astrocytes during fluorescence-activated cell sorting (FACS). Astrocytes were sorted based on ACSA2-positive, DAPI-positive (to exclude damaged cells), and tomato-negative (to exclude Cx3cr1-expressing cells) signals. ACSA2-positive cells were isolated from whole brains of control, cuprizone, or remyelinated conditions and subjected to single-cell sequencing (Fig. 4A). Non-astrocytic cells and doublets were removed from the data before downstream analysis.

**Figure 4.**
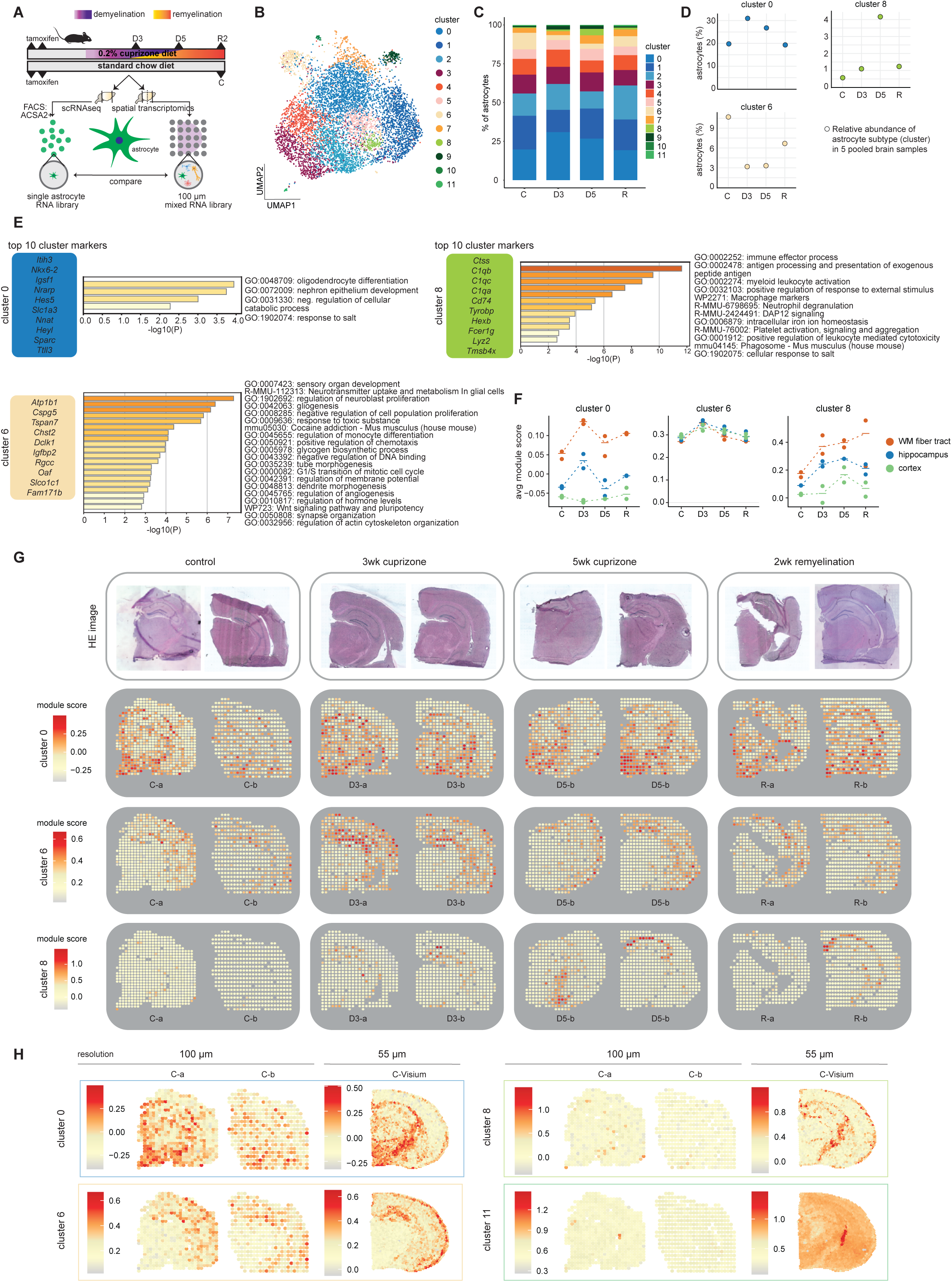
Regional heterogeneity of astrocytes in the mouse brain. **A**). Schematic experimental overview. Mice received either a cuprizone diet or a chow diet. Mice were sacrificed at 3 weeks (D3) or 5 weeks (D5) demyelination, or after a 2 weeks recovery period (R) (black arrowheads). Brain samples were further processed for spatial transcriptomics or single-cell sequencing. ACSA2-positive cells were sorted and profiled using single-cell RNA sequencing with the 10X Genomics platform. For single-cell RNA libraries one sample was sequenced where a sample represented 5 pooled mouse brains. Two mouse brains were used for ST. **B).** UMAP depicting the astrocyte clusters. Dots represent single cells, colors represent clusters. **C).** Stacked barplot of the relative distribution of astrocyte clusters (%) per experimental group. **D).** The relative abundance (%) of astrocyte clusters 0, 6, and 8 per experimental group as dot plot. One data point represents 5 whole brains pooled for single-cell RNA sequencing. **E).** The top 5 cluster markers for astrocyte clusters 0, 6, and 8 and the associated gene ontology terms. GO terms were retrieved from the Metascape database. **F).** Average module score for clusters 0, 6, and 8-associated markers in spatial spots of a given brain region. Data points represent ST mouse brain samples, colors indicate the brain region. **G).** Markers associated with astrocyte clusters 0, 6, and 8 are visualized as gene module scores in spatial spots. The color indicates the gene module score. **H).** Spatial plots of clusters 0, 6, and 8 of control mouse hemispheres using spatial transcriptomics with a resolution of 100 µm (left) and Visium with a resolution of 55 µm (right). The color indicates the gene module score per spatial spot. C = control, D3 = 3 weeks of cuprizone treatment, D5 = 5 weeks of cuprizone treatment, R = 2 weeks remyelination

After dimensionality reduction and clustering, 12 clusters of ACSA2-positive astrocytes were identified (Fig. 4B). Next, per cluster the differentially expressed genes (‘cluster markers’) and the associated GO-terms were determined (Table S6, Fig. S5A). ScCODA with adjustments for the one-sample scenario was used to identify credible changes in astrocyte subpopulation composition in demyelinated and remyelinated conditions. We did not identify any significant differences, but we used the scCODA-inferred log2 fold changes to prioritize and rank the largest differences in cluster composition (Table S6). Next, we investigated the spatial distribution of astrocyte subpopulations by visualizing and averaging gene module scores for cluster markers in ST data (Fig. S6).

The largest increase in relative abundance was detected within astrocyte cluster 8 (log2 fold change = 1.3) in 5-week cuprizone-treated mice compared to controls (Fig. 4C-D, Table S6). Astrocytes cluster 8 expressed complement genes (*C1qb, C1qc, C1qa),* and this cluster was associated with the GO terms “immune effector process”, “antigen processing and presentation” and “myeloid leukocyte activation” (Fig. 4E). Investigating the markers for astrocytes cluster 8, spatial plots suggested that these genes were mostly expressed in the hippocampus and WM fiber tract (Fig. 4F-G). Together, this suggested an immune-mediatory role for the astrocyte cluster 8 in the hippocampus and WM fiber tract that was most strongly associated with 5-week cuprizone-treated mice.

A subtle increase in relative abundance was detected in astrocyte cluster 0 for 3-week cuprizone-treated mice compared to controls (log2 fold change = 0.4) (Fig. 4C-D, Table S6). This astrocyte cluster had enriched expression of *Itih3, Hes5, Nrarp, Heyl, Slc1a3,* and *Sparc* (Table S5). The increased relative abundance of cluster 0 was in line with the previous observation of increased *Slc1a3* expression in the WM fiber tract of 3-week cuprizone-treated mice compared to controls in ST sections (Fig. 3B). For astrocyte cluster 0, relative abundance returned to control levels at 5-week cuprizone-treated and 2-week remyelinated conditions (Fig. 4D). Additionally, markers for astrocyte cluster 0 included *Cnp*, *Nkx6-2*, and *Hes5*, which were associated with the GO term “oligodendrocyte differentiation” (Fig. 4E). In the ST data, markers of astrocytes cluster 0 were expressed throughout multiple brain regions and were more enriched in the WM fiber tract and the hypothalamus compared to the cortex (Fig. 4F-G). In the WM fiber tract and hippocampus, an increased module score for astrocyte subpopulation 0-associated genes was observed at the 3-week cuprizone treatment. However, in the cortex, there were no discernible changes in response to the cuprizone treatment (see Fig. 4F-G). Possibly, astrocyte subpopulation 0 was less responsive to cuprizone treatment in the cortex than in other brain regions such as the hippocampus and WM fiber tract. In short, astrocyte cluster 0 was detected in multiple brain regions and showed a small increase in relative abundance in 3-week cuprizone-treated mice in whole-brain single-cell data. ST data suggested that cluster 0 was differently affected by the cuprizone diet across brain regions.

Lastly, astrocyte cluster 6 was depleted in 3-week (log2 fold change = –1.0) and 5-week (log2 fold change = –0.9) cuprizone-treated mice compared to controls (Fig. 4C-D, Table S6). This astrocyte cluster had enriched expression of *Atp1b1, Cspg5, Tspan7,* and *Chst2* compared to other astrocytes (Fig. 4E, Table S5). Cluster markers were associated with the REACTOME pathway “Neurotransmitter uptake and metabolism in glial cells”, and GO terms “synapse organization” and “regulation of the membrane potential”, which suggest a more GM-related functionality (Fig. 4E). In ST data, expression of astrocyte cluster 6 markers was enriched in the hippocampus and L1 of the cortex (Fig. 4F-G). Unexpectedly, the increase in astrocyte cluster 6 scores in ST data did not align with the decrease in the relative abundance of cluster 6 in single-cell data (Fig 4C-D, F-G). Possibly, this was due to brain region composition differences across ST sections: all ST samples at D3 contained complete hippocampal and L1 cortical structures, whereas these regions were not always complete at C, D5, and R samples. In contrast, for the single-cell data, a consistent brain region composition was expected for all samples. In summary, astrocyte cluster 6 was most prominently detected in the hippocampus and L1 cortical layer and showed a reduction in relative abundance in cuprizone-treated mice.

Higher spatial resolution is crucial to discern more specific regional astrocyte subpopulations To assess the spatial localization of the identified astrocyte subpopulations in the control mouse brain, we interrogated publicly available Visium spatial transcriptomics data (Fig. 4H). High-resolution Visium plots demonstrated regional diversity in astrocytes in the control mouse brain using smaller spatial spots (Fig. 4H). Comparing our astrocyte subpopulations in high-resolution Visium plots, a differential spatial distribution of astrocyte clusters 6 and 8 within the hippocampus region became visible in the control mouse brain (Fig. 4H, C-Visium). Most likely, cluster 6 reflects neuron-enriched hippocampal areas, and cluster 8 WM-fiber-enriched areas within the hippocampus.

The high-resolution Visium data also allowed for the annotation of an additional cluster. Cells in cluster 11 were thus far unannotated, since they did not express astrocyte-related marker genes (*Slc1a2, Slc1a3, Aldh1a1, Aqp4, Gfap*), but also did not clearly express markers specific to other cell types. In high-resolution Visium data, we observed that cluster 11 markers were highly abundant in regions around the choroid plexus (Fig. 4H, Fig. S5B). Congruently, the most enriched marker gene for cluster 11 (average log2 fold change = 7) was *Ttr* (Table S6), encoding a plasma transport protein with reported localization in the cytoplasm of choroid plexus-derived epithelial cells (Benson et al., 2010). Other marker genes for cluster 11 were *Enpp2, Krt18,* and *Kcnj13* (Table S5), which are reported markers for mouse epithelial cells in the choroid plexus (Dani et al., 2021). Together, this indicated that cluster 11 represented a small population of choroid plexus-derived epithelial cells. Comparing the ST and Visium platforms, the enhanced regional heterogeneity of astrocyte clusters 6 and 8, along with the specific localization of cluster 11 to the choroid plexus, were only detected with increased resolution (55 µm spots) (Fig. 4H). This highlights the importance of employing spatial technologies with higher resolution for a more fine-grained assessment of regional differences, cellular subpopulations, and transcriptional states.

## Discussion

In this study, we applied ST and scRNAseq to brain tissue from cuprizone-treated mice. We aimed to 1) identify gene expression differences during de– and remyelination, 2) examine regional gene expression alterations of glial cell type markers during de– and remyelination, and 3) assess the effects of cuprizone treatment on different brain regions. Using ST, we identified 70 DEGs within the WM fiber tract, indicating that astrocytes were important contributors to gene expression changes at the demyelination phase. On protein and gene expression levels, expression of GFAP was more affected in the cortex than in the corpus callosum following cuprizone treatment, an observation in line with the literature (Buschmann et al., 2012; Castillo-Rodriguez et al., 2022). By examining the expression of glial cell type markers, we observed a decreased expression of the mature OL gene *Mal* during demyelination, which was accompanied by an increase in the expression of reactive astrocyte markers in the hippocampus and WM fiber tract. At the remyelination phase, *Mal* expression returned to control levels, while astrocyte activation persisted. Combining scRNAseq of astrocytes with the ST data indicated that cuprizone-induced demyelination and remyelination differently affected astrocyte subpopulations across brain regions.

### Evaluating the ST platform

As an initial step, we aimed to validate if demyelination and remyelination could be detected in the cuprizone mouse model using ST. In this experiment, we applied ST on one hemisphere, meaning that both white and grey matter (GM) were included for analysis. We detected a decrease in the level of myelin genes at 3-weeks of demyelination (e.g. *Mog, Mal*), mostly in the WM fiber tract, indicating that ST with a resolution of 100 µm was sensitive enough to detect differences in gene expression related to demyelination. However, in ST data, most variation was driven by regional variation (WM, GM) and not by the cuprizone model itself, thereby hampering the detection of injury-induced changes. To overcome this regional heterogeneity between samples, we implemented multiple methods for annotation of brain regions and focused on cuprizone-induced gene expression changes within brain regions. In the future, region annotation methods could further improve by incorporating both the histology (HE image) and region-specific gene expression features to train machine learning models. This approach could aid the development of novel annotation tools that are objective and resistant to the influence of pathological and injury-induced gene expression variations.

### Temporal and regional responses of glial cells in the cuprizone model

Next, we studied the temporal and regional cuprizone effects on selected glial markers, to examine differences in the dynamics of glial cell responses. Using ST, we detected very low gene activity for low abundant cell types, such as microglia and OPCs, hampering the examination of the dynamics of genes in these cell types. Given the limited spatial resolution of 100 µm and spot-to-spot distance of 200 µm, ST might not be optimal for accurately capturing and characterizing microglia and OPCs, which are known to have relatively sparse representation in the brain. Moreover, microglia are relatively small cells, further hampering their detection using this ST platform.

Oligodendrocyte markers *Mal, Mbp, and Plp 1*showed a clear decrease in expression at 3-weeks of demyelination for the WM fiber tract, hippocampus, and cortex, which thereafter increased again. During demyelination (starting from week 3), compensatory mechanisms come into play to counteract the loss of mature OLs and myelin. At this stage, OPCs undergo proliferation, leading to an increase in *Plp1* expression (a marker for OPCs, immature and mature OLs), followed by an increase in *Mbp* expression (a marker for immature and mature OLs), as shown by the increase of these marker genes from D3 onwards.

Gene expression of reactive astrocyte markers *Gfap* and *Vim* peaked at 3-week demyelination in the WM fiber tract. At the protein level, GFAP was most abundant at 5-week demyelination in the corpus callosum, which was in line with previous literature (Buschmann et al., 2012; Castillo-Rodriguez et al., 2022). As reported before (Castillo-Rodriguez et al., 2022; Hibbits et al., 2012), *Gfap* gene expression decreased after 3-weeks of demyelination but did not return to control levels, indicating ongoing astrogliosis during 5-weeks of demyelination and 2-weeks remyelination. Loss-of-function studies investigating GFAP+ astrocytes have shown these astrocytes are required for maintaining myelin formation and OL maturation (Skripuletz et al., 2013; Tognatta et al., 2020). Ablation of GFAP+ astrocytes resulted in demyelination and myelin decompaction (Skripuletz et al., 2013; Tognatta et al., 2020). Therefore, persistent astrogliosis during the remyelination phase might be vital for the restoration of myelin and OL function.

### Astrocyte-related gene expression changes in the cuprizone model

When performing analysis on the WM fiber tract only, we identified astrocyte-related genes to be differentially expressed in 3-weeks of demyelination compared to the control. Genes such as *Clu* and *Slc1a3* were identified as DEGs in the WM fiber tract after cuprizone treatment. *Clu* encodes clusterin which is involved in the regulation of apoptosis and complement signaling, and enhanced levels of this protein were previously detected in astrocytes in WM lesions from MS donors. Depending on the astrocytic state (homeostatic vs. pathological), *Clu* can be secreted, or localized at the cytoplasm, or nucleus. In its secreted form, *Clu* serves as a complement inhibitor. When located in the nucleus, *Clu* has a pro-apoptotic effect on the astrocyte. On the contrary, when located in the cytoplasm, it functions as an apoptosis inhibitor (van Luijn et al., 2016). These multiple functions and states make it challenging to elucidate the role of *Clu* during de– and remyelination. *Slc1a3* encodes a gene for a glutamate transporter and upregulation of this gene is associated with excitotoxicity (Vallejo-Illarramendi et al., 2006). In MS, an increase in *Slc1a3* located in oligodendrocytes was observed in optic nerves (Vallejo-Illarramendi et al., 2006). Likely, increased expression of the glutamate transporter *Slc1a3* in cuprizone-fed mice could result in dysregulation of glutamate homeostasis in the brain, thereby contributing to neuronal dysfunction and degeneration.

To analyze the heterogeneity of astrocytes in the cuprizone mouse model, we performed scRNAseq and identified 12 astrocyte subpopulations. By combining these scRNAseq data with ST, we could predict in which brain regions these astrocyte subpopulations were enriched. Among the cuprizone-associated subpopulations, cluster 8 was enriched for expression of complement genes in the WM fiber tract and hippocampus (and not in L2-L6 of the cortex), indicating that cuprizone-induced neuroinflammation was more pronounced in WM areas, and less in GM areas. Markers of astrocyte cluster 8 were associated with the GO term “myeloid leukocyte activation”, suggesting possible astrocyte-microglia-immune cell crosstalk by this astrocyte subpopulation. Astrocytes are known to recruit microglia to the site of demyelination to clear myelin debris and if this process is disturbed, subsequent repair mechanisms are delayed (Skripuletz et al., 2013). Future studies investigating cell-cell interactions could demonstrate if this astrocyte subpopulation identified in cluster 8 is indeed involved in astrocyte-microglia communication. Astrocyte cluster 6 was identified in GM brain regions and showed a decrease in relative abundance under cuprizone conditions. Both astrocyte subpopulations also showed distinct expression patterns within the hippocampus itself, with cluster 6 prevalent in neuron-enriched areas and cluster 8 most abundant within WM-fiber-enriched hippocampal areas. For future studies, this stresses the importance of analyzing intra– and interregional astrocyte subpopulations separately in the context of remyelination failure and/or success.

Limitations of this study were the variation in brain region composition between some of the samples and the resolution of the ST spatial spots (100 µm) used here. Therefore, we validated part of our results in a publicly available Visium ST data set (55 µm spatial spots) and used our cuprizone scRNAseq data to further investigate the astrocyte-related changes in different brain regions. At 55 µm resolution, we were able to map the heterogeneity of astrocytes in the control mouse brain more precisely, whereas this was more challenging at 100 µm resolution. This comparison underscores the importance of increased resolution in ST technologies. Furthermore, using ST it was not possible to determine whether the detected changes across the experimental groups were due to variations in the abundance of different cell types (e.g. increased GFAP+ astrocyte numbers) or if the changes in gene expression occurred within specific cell populations (e.g. increased GFAP expression within astrocytes). To address this issue, future studies would require spatial analysis at a cellular resolution (Stereo-seq (Xia et al., 2022), CosMx^TM^, Xenium analyzer, or a forthcoming Visium HD from 10X Genomics).

## Conclusion

In conclusion, ST was sensitive enough to detect demyelination processes and enabled unbiased characterization of multiple brain regions in a single tissue section. This facilitated the identification of region-associated changes in gene expression following cuprizone treatment. The main limitation encountered using ST is that the lack of a single-cell resolution hindered definite conclusions on less abundant cell types, such as OPCs and microglia, and that possible disease-driven alterations in cell type composition were potentially masking cell type-specific gene expression changes. ST approaches at (near) single-cell resolution in combination with computational deconvolution and segmentation methods are required to delineate the role of specific cellular subpopulations in animal models or diseases. We identified astrocyte heterogeneity across brain regions of control and cuprizone-treated mice. Cuprizone-induced demyelination changed the expression of astrocyte subpopulation-associated genes differently across the WM fiber tract, hippocampus, and cortex. The effect of cuprizone treatment on astrocyte cluster 0, which was associated with oligodendrocyte differentiation, and cluster 8, which was associated with immune-related processes, seemed to depend on the brain region. The characterization of (intra)regional astrocyte subpopulations could aid the development of novel strategies to target demyelination and remyelination processes and to manipulate their involvement in diseases such as MS.

## Acknowledgments

We would like to thank Geert Mesander, Johan Teunis en Theo Bijlsma from the flow cytometry unit of the UMCG for sorting the ACSA2+ positive cells and Klaas Sjollema from the UMCG Imaging and Microscopy Center (UMIC) for the support when imaging tissue sections with the TissueFaxs and Zeiss Cell Discoverer 7. We thank the Research Sequencing Facility of the UMCG, in particular Diana Spierings, for sequencing and discussion of ST data. We thank Zaneta Andrusivova, Michaela Asp, and Annelie Mollbrink from the SciLifeLab at Karolinska Institutet, Sweden for their help and hospitality during workshops on the spatial transcriptomics technique.

## Ethics Approval

All animal experiments were approved by the national central authority for scientific procedures on animals (CCD) and performed in accordance with ethical regulations (Permit# AVD105002015360).

## Authors’ contributions

AM and MHCW performed the experimental work. MM and EMW supported the experimental work. AMA, EG, and MK performed the data analysis. WB, BJLE, and SMK conceptualized and supervised the project. AM, MHCW, and AMA wrote the manuscript. All authors read and contributed to the manuscript.

## Data availability

The spatial transcriptomics sequencing datasets generated for this study have been deposited in the Synapse database under accession number syn52597881. Foundational data for bar graphs is available in the Supplementary Tables.

## Code availability

The code that supports the findings of this study is available from the corresponding authors upon request.

## Figure legends

**Supplemental figure 1.**
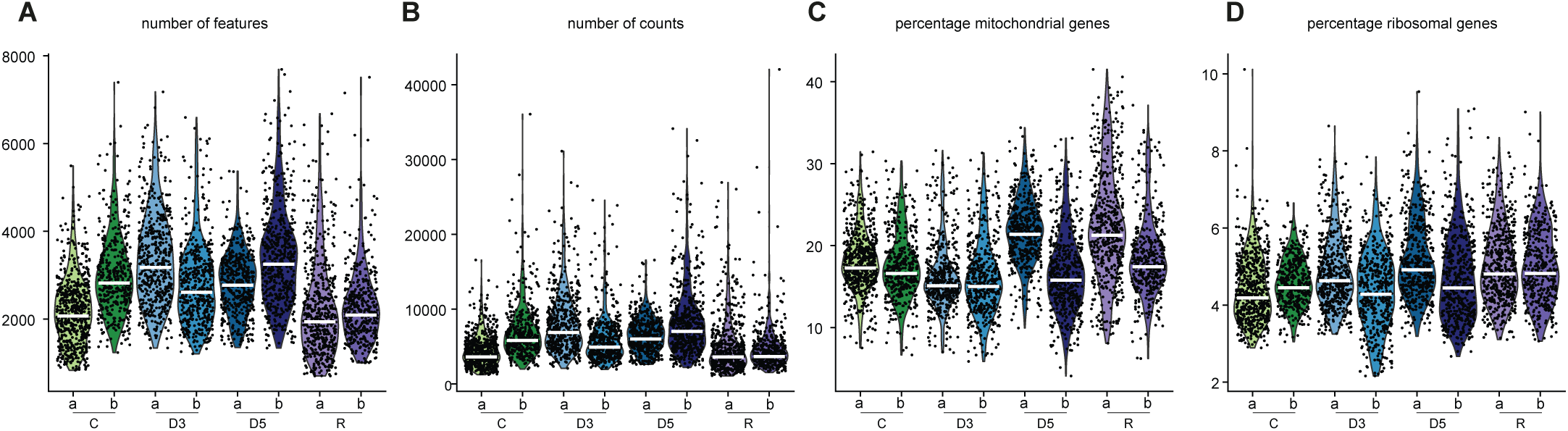
Quality control of 100µm-resolution spatial transcriptomic spots in mouse brain samples. **A**). Quality control parameters of the number of features **B).** number of UMI counts **C).** percentage of mitochondrial genes and **D).** percentage of ribosomal genes depicted in violin plots for the individual samples per experimental group. The horizontal line of the violin plots depicts the median. C = control, D3 = 3 weeks of cuprizone treatment, D5 = 5 weeks of cuprizone treatment, R = 2 weeks remyelination

**Supplemental figure 2.**
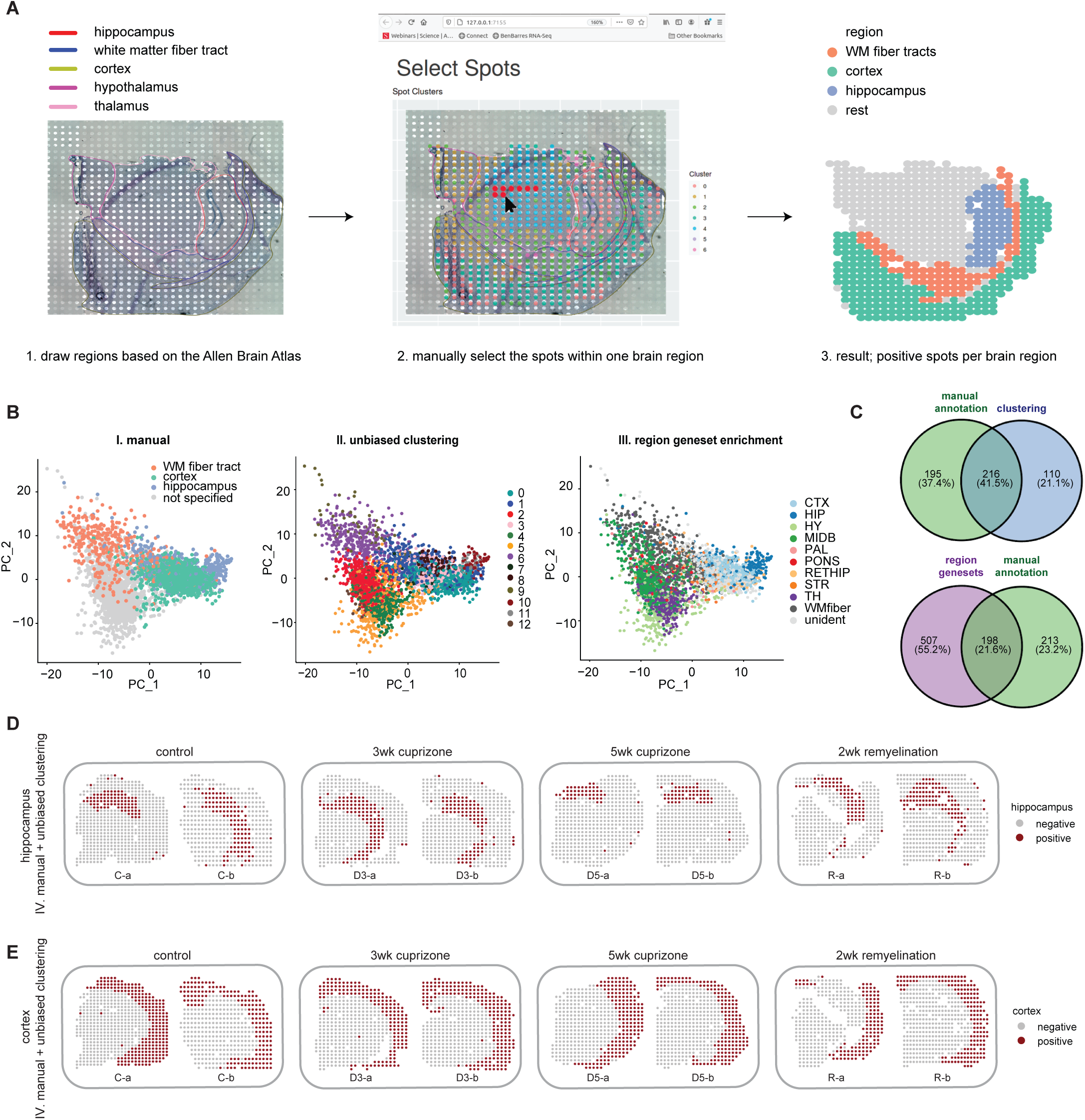
Comparison of region annotation methods. **A**). Schematic representation of the manual annotation method. **B).** PCA plots of spatial spots with colors indicating annotation by the manual method (I), unbiased clustering method (II), and region geneset enrichment method (III). **C)** For the WM fiber tract, the upper Venn diagram depicts the overlap between the manual annotation and the clustering-based annotation method. The bottom Venn diagram depicts the overlap between the manual and the gene set enrichment annotation method. **D).** Annotation results from the combinatorial approach (IV) of the hippocampus. Dark red spots indicate the spots annotated as hippocampus. **E).** Results of the combinatorial approach (IV) of the cortex. Dark red spots indicate the spots annotated as cortex. C = control, D3 = 3 weeks of cuprizone treatment, D5 = 5 weeks of cuprizone treatment, R = 2 weeks remyelination

**Supplemental figure 3.**
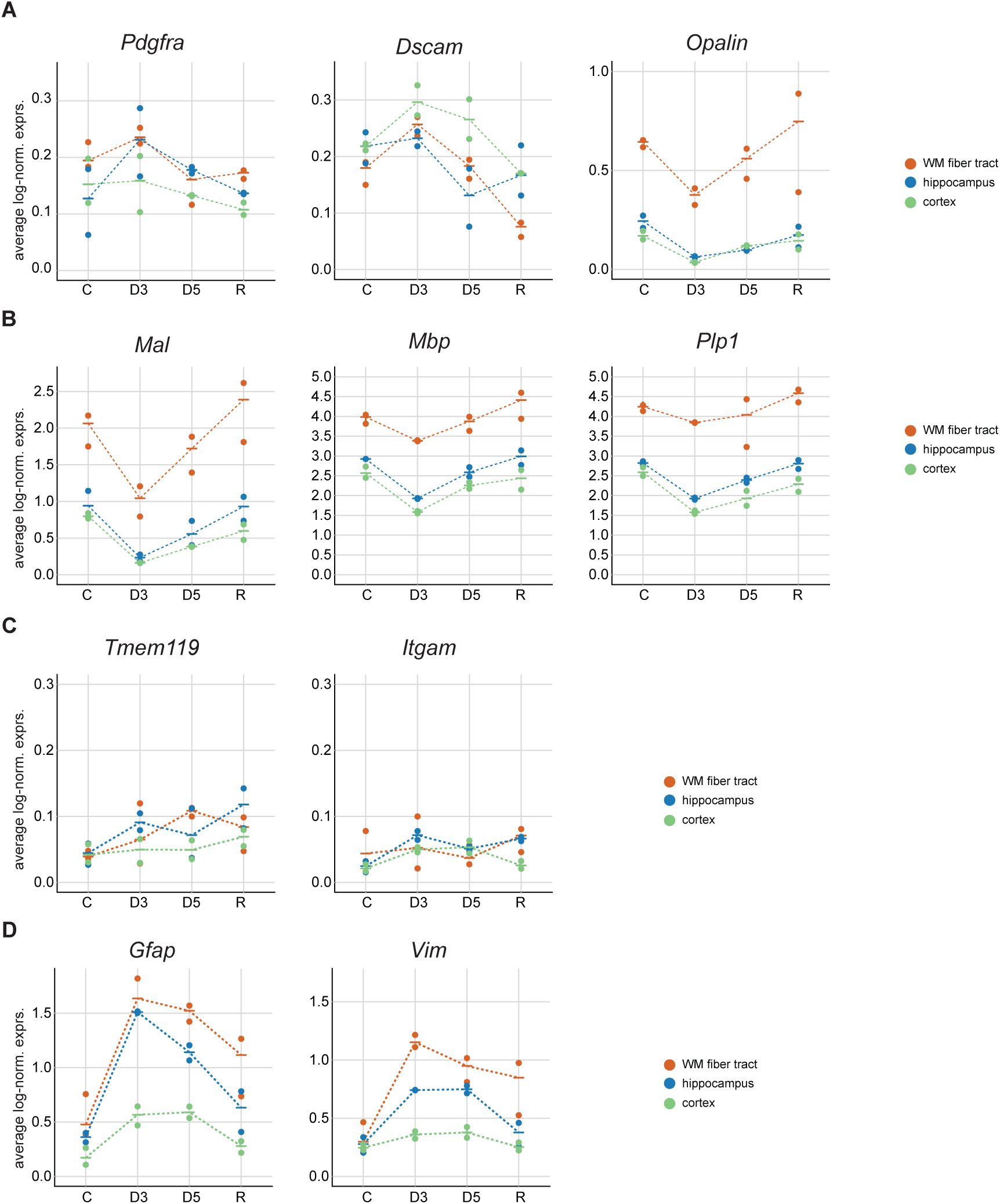
Selected transcriptional markers for oligodendrocyte lineage cells (A-B), microglia (C), and astrocytes (D). Data points represent mouse brain samples, colors indicate the brain region. The horizontal continuous line represents the averaged normalized expression of spatial spots in a given region across mice. Normalized gene expression counts were obtained using a natural logarithm. C = control, D3 = 3 weeks of cuprizone treatment, D5 = 5 weeks of cuprizone treatment, R = 2 weeks remyelination

**Supplemental figure 4.**
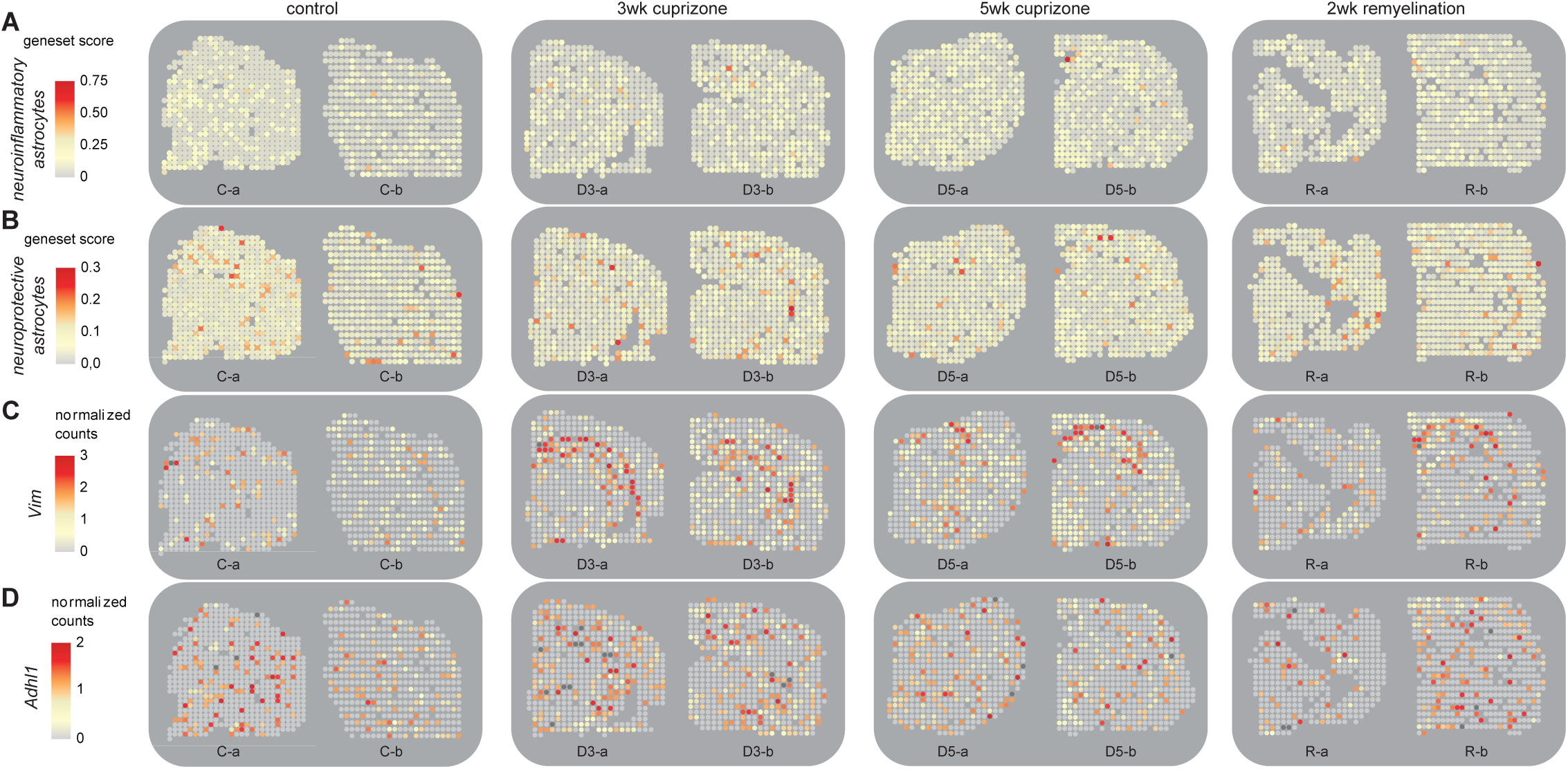
Expression of astrocyte genes in cuprizone-treated mice. A-B). Representative spatial plots depicting geneset scores of genes associated with neuroinflammatory **(A)** and neuroprotective **(B)** astrocytes (Liddelow et al., 2017) per experimental group. The color bar indicates gene set module scores. **C-D).** Representative spatial gene expression plots of *Vim* **(C)** and *Aldh1l1* **(D)** per experimental group. The color bar indicates log-normalized gene expression level. C = control, D3 = 3 weeks of cuprizone treatment, D5 = 5 weeks of cuprizone treatment, R = 2 weeks remyelination

**Supplemental figure 5.**
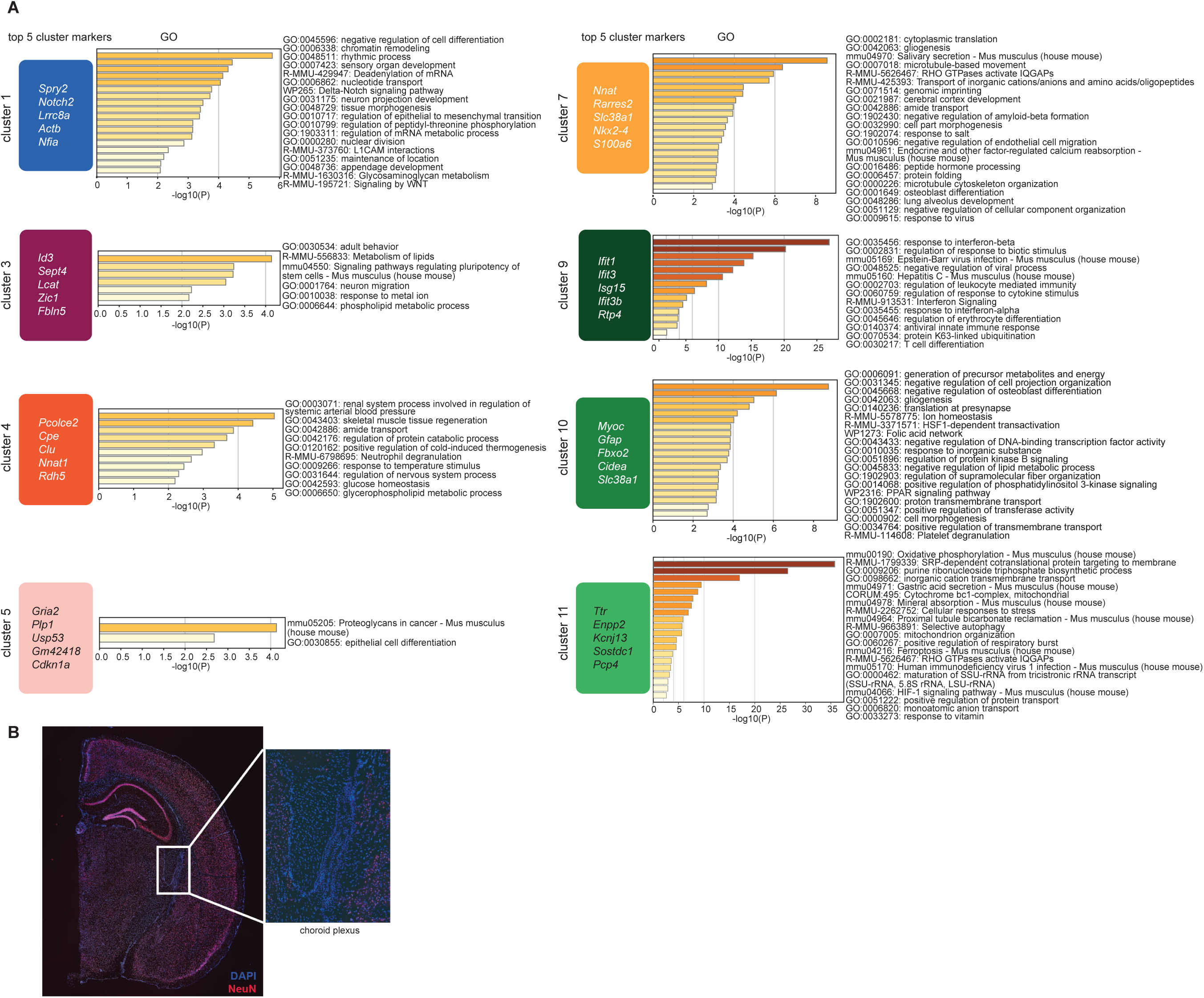
Gene ontology enrichment per astrocyte cluster detected with single-cell RNA sequencing. **A**). The top 5 cluster markers depicted per astrocyte cluster were ranked based on the highest log-fold change. Bars represent the significance of the cluster marker-associated gene ontology term retrieved from the Metascape database. **B).** Fluorescent image of a fresh-frozen coronal adult mouse brain section stained for NeuN (red) and DAPI (blue). Data was derived from a publicly available dataset from 10X Genomics. The inset shows the area of interest for the cluster markers of astrocyte subpopulation 11.

**Supplemental figure 6.**
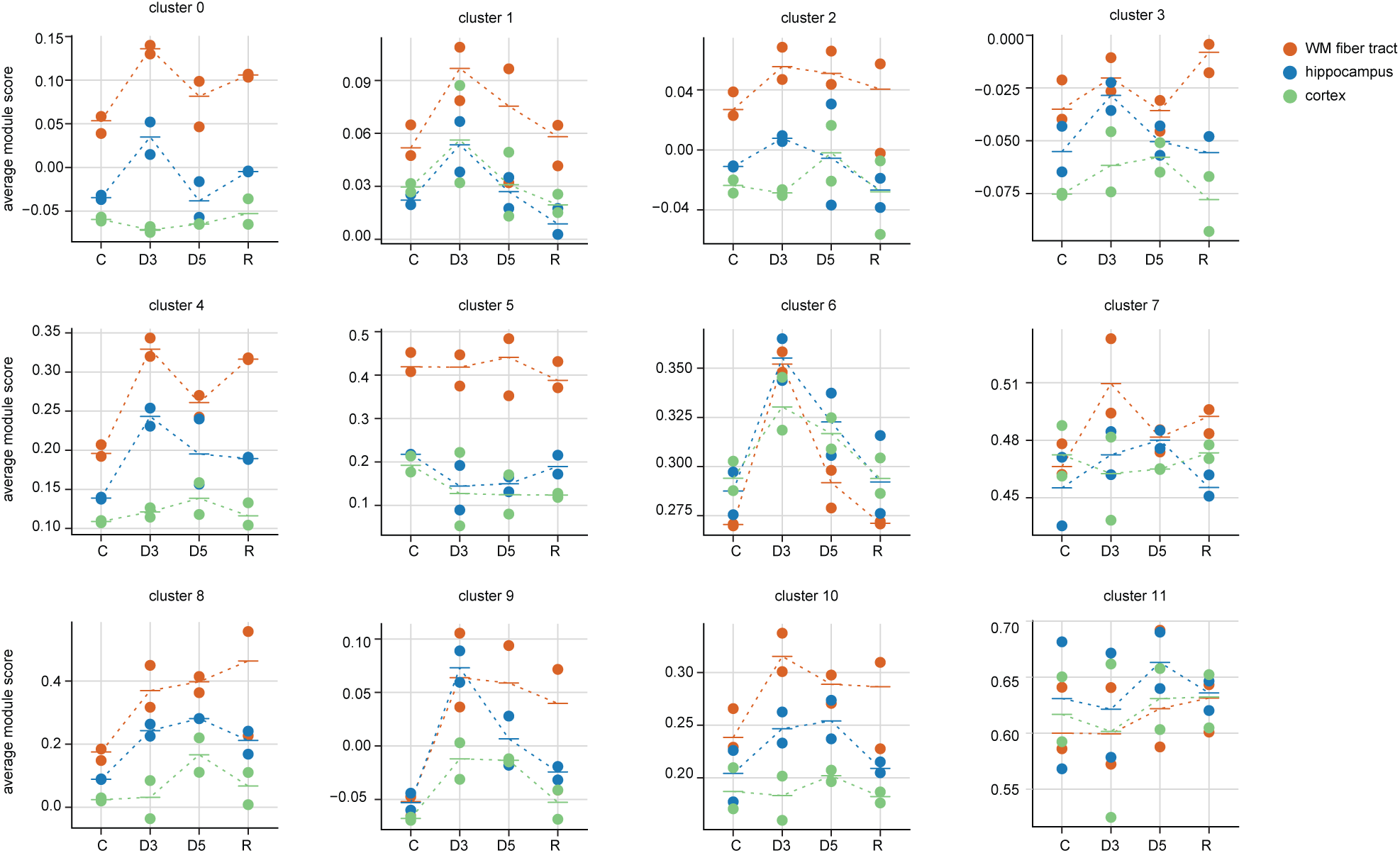
Average module scores of astrocyte cluster markers in the WM fiber tract, hippocampus, and cortex. Data points represent mouse brain samples, colors indicate the brain region. The horizontal continuous line represents the average module score of spatial spots in a given region across mice. Module scores were computed for transcriptional markers associated with the astrocyte clusters in a given brain region. C = control, D3 = 3 weeks of cuprizone treatment, D5 = 5 weeks of cuprizone treatment, R = 2 weeks remyelination.

## Supplementary tables

Excel file containing tables S1-S9

**Table S1:** sample information of the acute cuprizone mouse model (ST platform)

**Table S2:** region-specific genesets for mouse brain region annotation for method III: gene set enrichment, and A1, A2, PAN-reactive genesets for spatial plots (ST platform)

**Table S3:** differentially expressed genes within the WM tract in the acute cuprizone mouse model (ST platform)

**Table S4:** cell type gene set enrichment for differentially expressed genes within the WM fiber tract

**Table S5:** GFAP protein levels in the corpus callosum and cortex and statistics corresponding to Figure 3H

**Table S6:** sample and sequencing information for the single-cell RNA sequencing experiment using the cuprizone mouse model (10X Genomics platform)

**Table S7:** cluster markers per astrocyte cluster (10X Genomics platform)

**Table S8:** single astrocyte cluster distribution per experimental group and scCODA results (10X Genomics platform)

**Table S9:** GO terms associated with astrocyte cluster markers (10X Genomics platform)

